# Internet-connected cortical organoids for project-based stem cell and neuroscience education

**DOI:** 10.1101/2023.07.13.546418

**Authors:** Matthew A.T. Elliott, Hunter E. Schweiger, Ash Robbins, Samira Vera-Choqqueccota, Drew Ehrlich, Sebastian Hernandez, Kateryna Voitiuk, Jinghui Geng, Jess L. Sevetson, Yohei M. Rosen, Mircea Teodorescu, Nico O. Wagner, David Haussler, Mohammed A. Mostajo-Radji

**Author notes:** These authors contributed equally to this work (Matthew A.T. Elliott, Hunter E. Schweiger, Ash Robbins).

## Abstract

The introduction of internet-connected technologies to the classroom has the potential to revolutionize STEM education by allowing students to perform experiments in complex models that are unattainable in traditional teaching laboratories. By connecting laboratory equipment to the cloud, we introduce students to experimentation in pluripotent stem cell-derived cortical organoids in two different settings: Using microscopy to monitor organoid growth in an introductory tissue culture course, and using high density multielectrode arrays to perform neuronal stimulation and recording in an advanced neuroscience mathematics course. We demonstrate that this approach develops interest in stem cell and neuroscience in the students of both courses. All together, we propose cloud technologies as an effective and scalable approach for complex project-based university training.

**HIGHLIGHTS:**

- Development of cortical organoids as pedagogical tools for undergraduate education.
- Organoids implemented in a tissue culture course through cloud-enabled microscopy.
- Multielectrode arrays allow for live organoid manipulation in a mathematics course.
- Students self-report increased interest in neuroscience and stem cells topics.

## INTRODUCTION

Pluripotent stem cell (PSCs)-derived 3D cortical organoids are transforming the study of human development, disease, and evolution (Mostajo-Radji et al., 2020; Nowakowski and Salama, 2022; Paulsen et al., 2022; Pollen et al., 2019; Velasco et al., 2020). Organoids model several aspects of corticogenesis, including the emergence of cell types, such as neuronal progenitors, projection neurons, interneurons, astrocytes, and other glia cell types (Nowakowski and Salama, 2022; Paulsen et al., 2022; Pollen et al., 2019; Velasco et al., 2020); as well as formation and maturation of neuronal networks (Cai et al., 2023; Fair et al., 2020; Sharf et al., 2022; Trujillo et al., 2019).

The use of cortical organoids in academic research and biotechnology has drastically increased in recent years (Salick et al., 2021). For example, cortical organoids are used as tools for drug discovery (Salick et al., 2021), to study genetic mutations underlying disease (Paulsen et al., 2022; Wang, 2018), and to study infectious diseases that affect the brain, such as Zika and COVID-19 (Andrews et al., 2022; Garcez et al., 2016). Given their applications and the trends in the use of these models in research and development, there is a growing need for students to be trained in stem cell culture and cortical organoid generation, manipulation, and analysis.

Tissue culture courses are challenging to implement due to the high costs of infrastructure and supplies associated with the course, as well as the danger of exposure of students to biohazardous materials (Bowey-Dellinger et al., 2017; Robinson et al., 2020). Indeed, the large majority of undergraduate biology students are not trained in cell culture during their education (Bowey-Dellinger et al., 2017). The students who are fortunate to learn tissue culture techniques are usually trained in either plant cell cultures (Harahap et al., 2019; Siritunga et al., 2012) or simple animal cell culture, such as growth and passage of established cell lines (Bowey-Dellinger et al., 2017; McIlrath et al., 2015; Mozdziak et al., 2004; Phelan and Szabo, 2019) or primary cells (Burdo, 2013; Catlin et al., 2016; Lemons, 2012; Haskew-Layton and Minkler, 2020). While remarkable, these approaches are not sufficient to familiarize students with increasingly complex models, such as PSC culture, differentiation, and formation of 3D models, including organoids.

Despite the high interest of students in the topic (Bajek and Drewa, 2011), classes that focus on stem cell use and applications are mostly theoretical (Ferreira et al., 2019; Perlin et al., 2013. Pierret and Friedrichsen, 2009; W Sandoval, 2022). Courses with a lab component usually take advantage of invertebrate organisms such as planaria worms (Accorsi et al., 2017; Ochoa et al., 2019) or hydra (Larouche et al., 2020). On the other hand, courses that integrate mammalian stem cell culture are limited to elementary models such as culture and rapid differentiation of primary multipotent stem cells derived from the bone marrow of rodents (Jin et al., 2018), not PSC-derived models such as complex tissues and organoids. These fail to meet the need for students trained in high-level biomedical cell models. Indeed, students are usually first exposed to PSC-derived models in special extracurricular undergraduate research experiences, such as internships (Halverson et al., 2009). These research experiences are disproportionately accessible to students from privileged backgrounds and institutions (Grineski et al., 2018). To create a diverse workforce in the stem cell field we must find alternative approaches that can effectively train underserved students (Hruska et al., 2022).

Internet-enabled technologies can be a powerful tool to overcome this gap. When laboratory equipment is connected to the cloud, scientists and students can easily access and manipulate these technologies remotely (Baudin et al., 2022a; Ly et al., 2021; Nagarathna et al., 2022; Parks et al., 2022). Moreover, this approach allows exceptional platform scalability, allowing dozens or hundreds of users simultaneously (Baudin et al., 2022b). In the classroom, internet-enabled technologies have been used to drive project-based learning (PBL) approaches in a variety of settings, including the use of microscopes for remote observations of behaviors of model organisms and cell cultures (Baudin et al., 2022b; Hossain et al., 2016; Ly et al., 2021); the use of sensors to monitor environmental conditions (Tabuenca et al., 2023; Tsybulsky and Sinai, 2022); and even the use of lab-on-a-chip microfluidic devices to perform bacterial detection in water samples (Sano et al., 2023).

Here, we take advantage of internet-enabled technologies to introduce cortical organoid PBL in two different university courses: an introductory tissue culture course, where students monitor the growth of organoids in the presence of different drugs; and an advanced neuroscience mathematics course where students stimulate and record from cortical organoids using multielectrode arrays (MEAs). In both approaches, the students can perform PBL over several days. We showed that internet-enabled PBL leads to a high level of interest in neuroscience and stem cell biology in the students in both courses. Altogether we propose that internet-enabled PBL can become an effective approach to deliver complex concepts in stem cell and organoid culture.

## RESULTS

### Internet-connected microscopy enables the incorporation of organoid modules in a tissue culture undergraduate course

PBL is an effective teaching approach for complex topics in biology, including teaching students who have traditionally underperformed in STEM (Baudin et al., 2022b; Ferreira et al., 2019; Petersen, 2021; Sano et al., 2023). We have previously shown that low-cost in-incubator internet-connected microscopes can perform longitudinal imaging of cortical organoids and allow for the systematic tracking of organoid size and morphology over time (Ly et al., 2021), making them ideal tools to perform remote PBL training in the classroom (Baudin et al., 2022b).

As a proof of principle, we integrated an organoid tracking module within the Techniques in Tissue Culture course at the University of San Francisco (USF), located at approximately 125 km from the experimental site at the University of California Santa Cruz (UCSC). This course has a strong hands-on component in which students learn basic techniques, including 2D culture of mammalian cancer cell lines. The students are exposed to the theory behind organoid generation, but the current curriculum does not have an organoid generation and culture experimental component. In this iteration of the course, 60% of the students were 3rd year undergraduate students, 30% of them were 4th year undergraduate students and 10% of them were students in their 5th year or above.

To design a project of interest to the students, the professor who was leading the course had a brainstorming session with the students ahead of the experiment. In this session the students were asked to nominate potential drugs that they thought would affect organoid growth. They nominated 2 drugs: staurosporine, a nonselective protein kinase inhibitor that is used in the research setting to induce apoptosis (Chae et al., 2000); and camptothecin, a selective DNA topoisomerase type I inhibitor that has been used to treat different types of cancers (Martino et al., 2017).

To accommodate the students’ requested project, we continuously imaged mouse cortical organoids starting at differentiation day 26 using a Streamscope, an in-incubator low-cost microscope that has the ability to both record image stills and video, which are then streamed over the internet through YouTube (Baudin et al., 2022b) (Figures 1A-B). The organoids were grown in the presence of either staurosporine or camptothecin. As control, organoids were grown without the presence of any drug. The Streamscope imaged organoids every 60 seconds for 3 days (Figure 1A). This approach allowed us to constantly produce data and promote student engagement. The students were divided into groups of 3-4 and were asked to measure the maximal organoid area over time. They were then asked to discuss the effects of the drugs as a class. The students observed a reduction in organoid size in organoids that were treated with staurosporine (Figure 1B). Consistent with the fact mouse cortical organoids of this age contain mostly postmitotic neurons (Eiraku et al., 2008; Kadoshima et al., 2013; Park et al., 2023), the students did not observe a reduction in organoid size in camptothecin-treated organoids (Figure 1C). We therefore show that internet-connected microscopes can effectively be used in the classroom setting to perform PBL approaches using organoid cultures.

**Figure 1.**
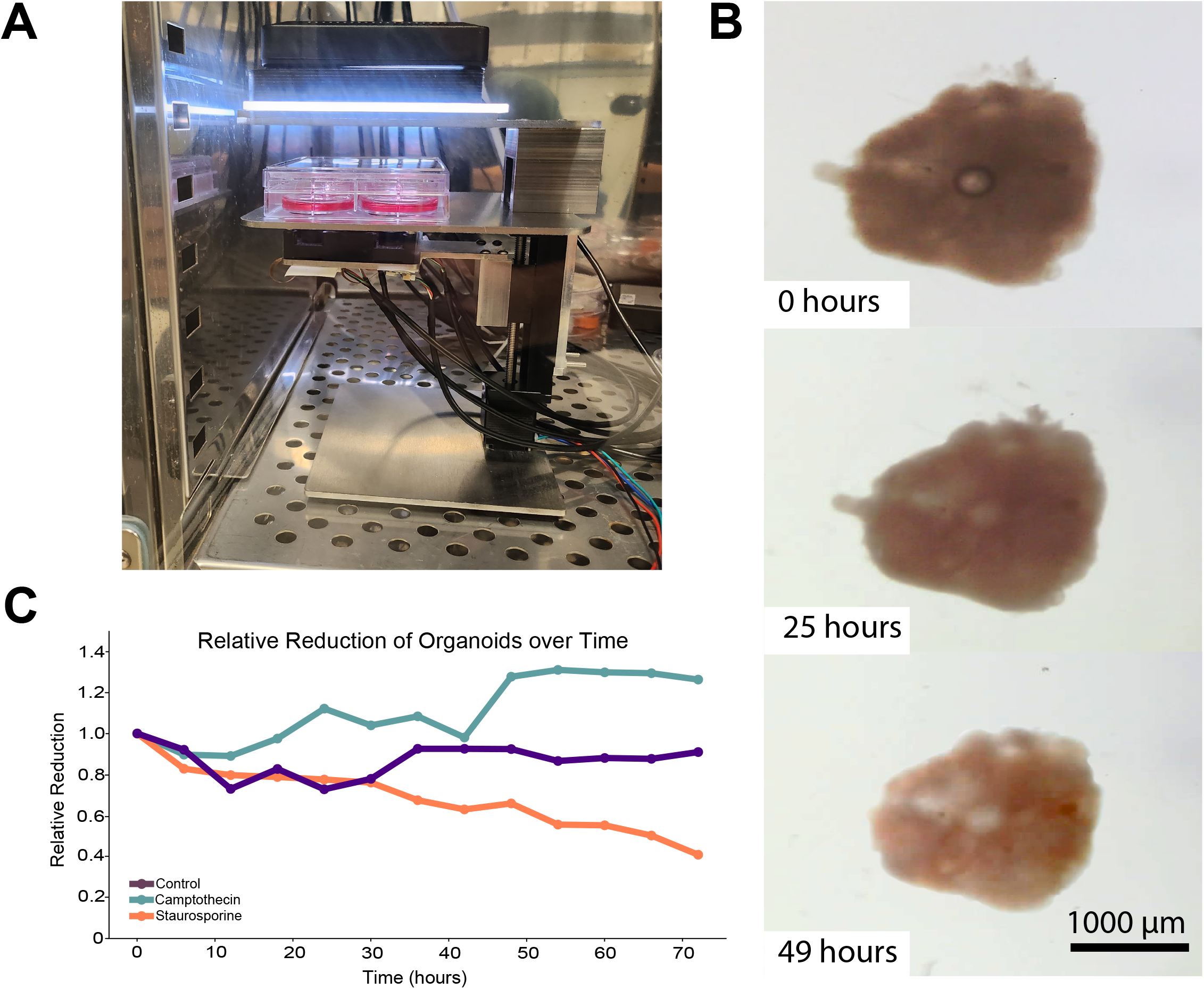
Internet-enabled microscopy enables longitudinal organoid tracking. A. Experimental setup showing a Streamscope inside a tissue culture incubator tracking 6 organoids. B. Example of an organoid exposed to the proapoptotic drug Staurosporine. C. Example results obtained from a group of students calculating the maximal organoid area over 72 hours. For this experiment, each group of students measured 3 individual organoids, one of each condition.

### Remote cortical organoid culture leads to strong interest in stem cell topics

After the completion of the course, we surveyed the students to understand their satisfaction with the technology and the course, as well as their interest in topics related to stem cells and organoids. All students who were part of the course responded to the survey (Figure 2).

**Figure 2.**
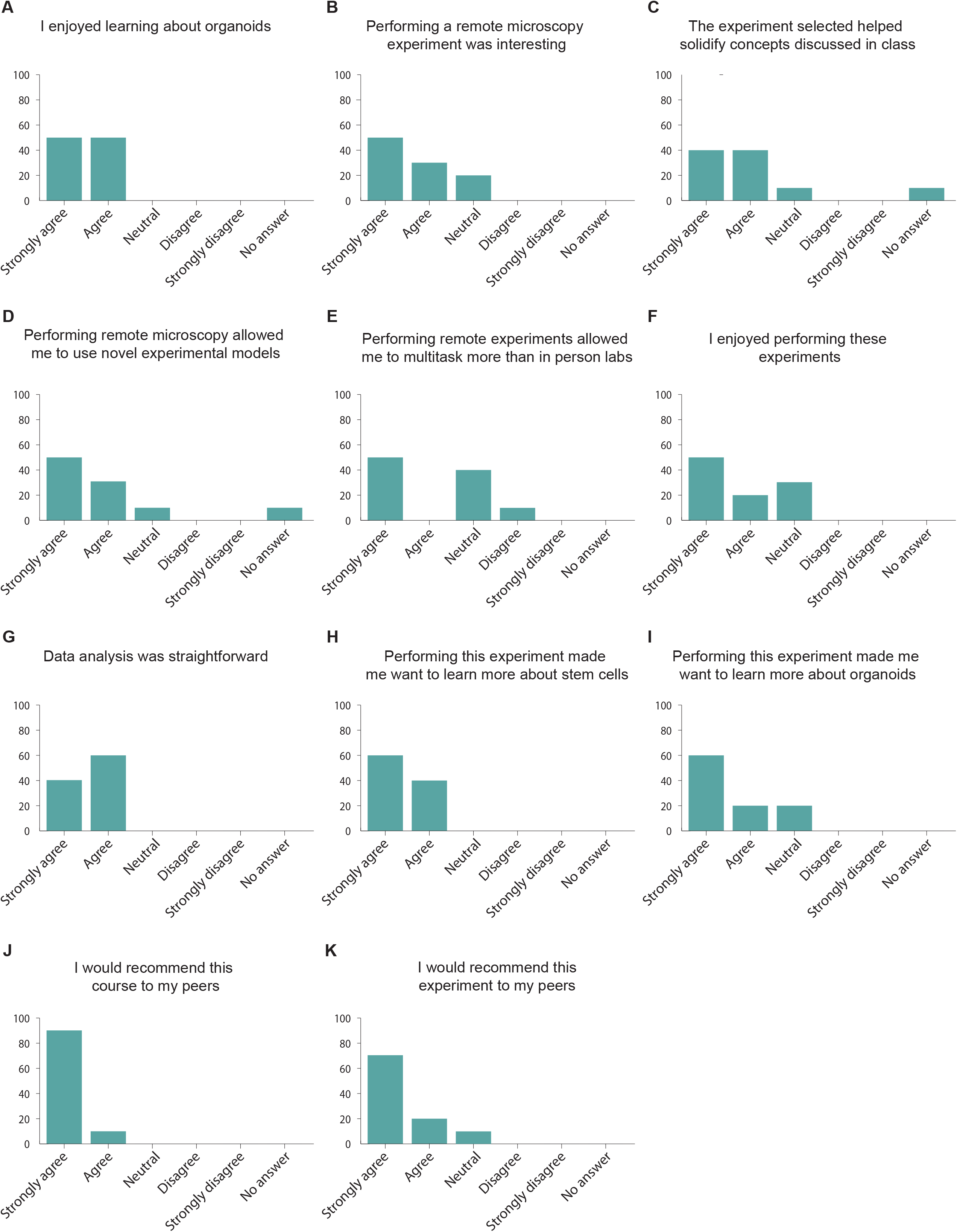
Students who perform remote cortical organoid culture report positive feelings and strong interest in stem cell topics. (A-K) Responses of the Techniques in Tissue Culture students to the post-experiment survey. n = 10 students.

To measure their satisfaction and interest, we asked the students their level of agreements with different statements. All students said that they enjoyed learning about organoids (Figure 2A), and 80% of them thought that performing these experiments helped them solidify the concepts learned in class (Figure 2B).

We then asked the students how they felt performing experiments using remote microscopy. We found that 80% of the students thought that performing remote microscopy experiments was interesting (Figure 2C) and that it allowed them to use novel experimental models (Figure 2D). Interestingly, only 50% of the students thought that they were able to multitask while performing these experiments (Figure 2E), suggesting that for at least half of the students, performing remote experiments requires them to concentrate to similar levels than in person laboratories. Nevertheless, 70% of the students reported that they enjoyed performing these experiments (Figure 2F) and all thought that data analysis was straightforward (Figure 2G).

When asked how this project affected their interest in learning about stem cells and organoids, all the students reported that this project made them want to learn more about stem cells (Figure 2H), while 80% of them reported that they want to learn more about organoids (Figure 2I). Finally, we find that 90% of the students would recommend this project to their peers (Figure 2J), while all students reported that they would recommend this class to others (Figure 2K). Altogether, we conclude that internet-enabled microscopy is an effective approach to introduce students to organoid culture, while increasing their interest in learning topics on stem cells and organoids.

### Electrophysiology software programs run on organoids in a neuroscience mathematics course

A landmark in neuronal maturation is the acquisition of electrophysiological properties (Fair et al., 2020). These properties follow a stereotypical cell-type specific identity (Cadwell et al., 2016; Gouwens et al., 2019; Ye et al., 2015; Zeng and Sanes, 2017). Given the great diversity of neurons in the cerebral cortex, it is expected that complex models, such as organoids, will have different electrophysiological profiles representative of their cell type composition (Passaro and Stice, 2021). Indeed, the emergence of complex networks in cortical organoids is an active area of research that is expected to continue growing over the next few years (Cai et al., 2023; Fair et al., 2020; Sharf et al., 2022; Trujillo et al., 2019).

One of the most exciting aspects of the emergence of neuronal circuits is that it provides scientists with the ability to study how neurons in organoids “learn” by adapting to stimuli provided to them (Zafeiriou et al., 2020). Neuronal plasticity can be evoked in cerebral organoids by sending electrophysiological stimulus patterns (Zafeiriou et al., 2020). These patterns are written using computer code. Therefore, an organoid’s neuronal circuitry can be “programmed” to produce different types of predictable neuronal responses.

To date, PBL-based teaching of electrophysiology concepts in the classroom have been limited to either simulation of single neurons using web-based apps (Cannon and Hammond, 2008; Demir, 2006; Formella-Zimmermann et al., 2022; Yamamoto et al., 2023) or the use of simple devices to activate and record neuromuscular electrophysiological signals in either invertebrates or the students’ own muscles (Ferreira et al., 2019; Hanzlick-Burton, et al., 2020; Marzullo and Gage, 2012; Ramadan and Ricoy, 2023). While remarkable, these experiments do not offer the capability to ask complex questions in circuit assembly and brain function. Specifically, students do not have the ability to program their own experiments, where they see how neuronal circuitry responds to their code. This is made possible using high-density multielectrode arrays (HD MEAs), in which thousands of electrodes can stimulate and record from a single circuit (Miccoli et al., 2019). For example, the MaxOne system from MaxWell Biosystems has been used to record circuit activity in cortical organoids through its 26,400 electrodes (Paulsen et al., 2022; Sharf et al., 2022). Yet, due to the high costs and experimental complexity associated with these experiments, HD MEAs have not been introduced in the classroom. We have previously developed an internet-connected electrophysiology platform that allows the users to record from MEAs remotely (Voitiuk et al., 2021). This technology could become key for cloud-enabled PBL (Baudin et al., 2022b), in which students use experimental platforms that are not located at or owned by their own institutions.

To deploy this technology for neuroscience education, we integrated it into the Mathematics of the Mind course at UCSC. This course is a mixed upper-level undergraduate and beginning graduate class where most enrolled students are from quantitative fields such as computer science, physics, applied mathematics, and engineering. Students were taught concepts and given supplementary readings within topics such as tetanus-induced potentiation dependent on connectivity (Jimbo et al., 1999), Hebbian theory (Hebb, 2005), spike-time dependent plasticity (Caporale and Dan, 2008) and homeostatic regulation (Harnack et al., 2015). For the students to properly construct experiments, they were introduced to applied electrophysiology techniques like spike sorting, a neurophysiology technique that allows neuronal spikes to be clustered based on similarities (Quiroga, 2012). To understand how performing live electrophysiological experiments impacted the students, we first surveyed them after the introduction of these concepts but before performing the experiments (Figure 3). Of the 24 students enrolled in the class, 18 responded to the precourse survey. We found that the large majority (83.7%) of the students had previous experience in mathematics and computer programming (Figures 3A and 3B). In contrast, 66.6% of the students reported having little to no previous stem cell biology experience (Figure 3C) and 94% of the students reported having little to no previous experience with neuroscience (Figure 3D).

**Figure 3.**
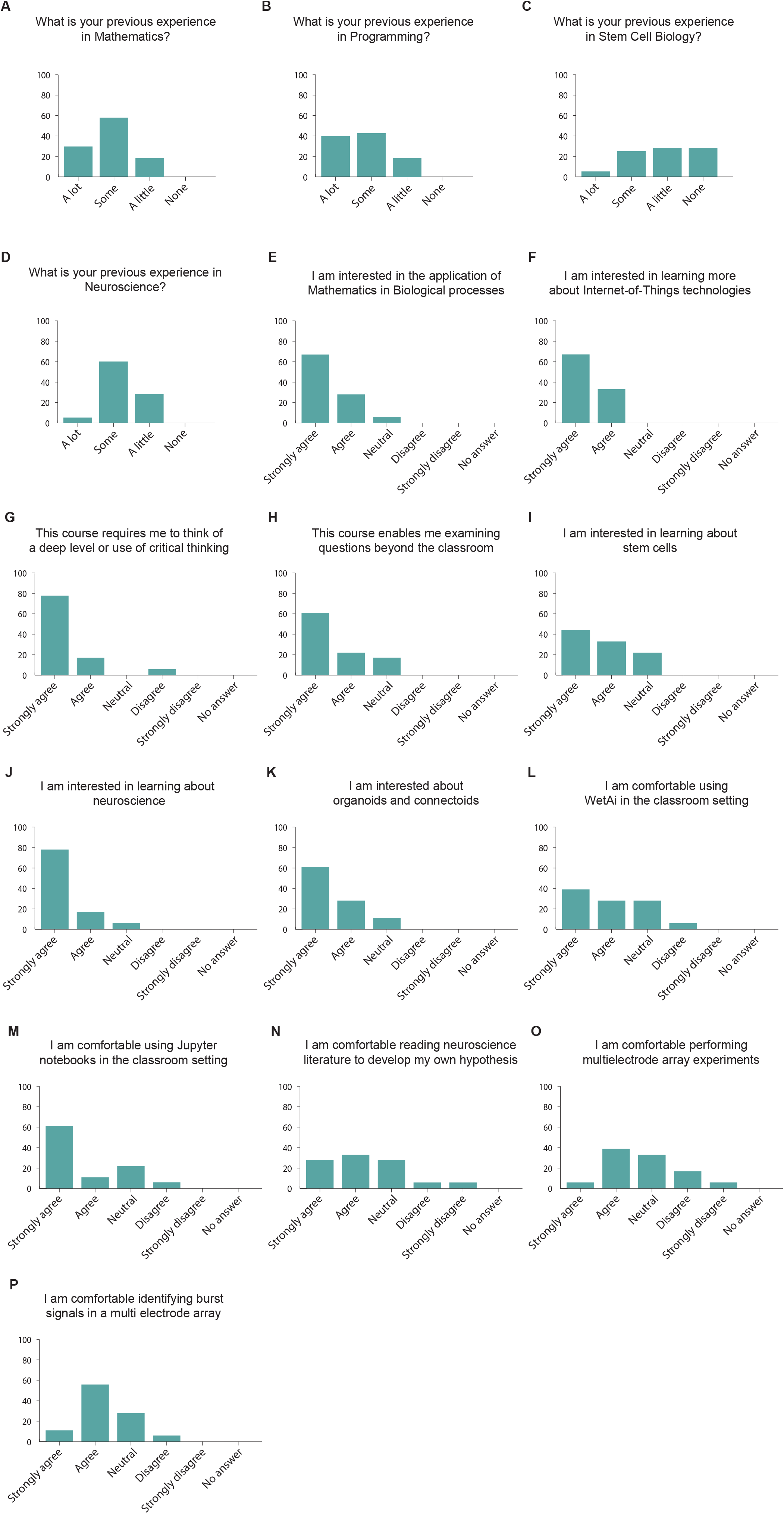
Students’ previous experience and reported interest in topics related to mathematics, neuroscience and stem cell biology. A-P. Students’ answers to a survey assessing their level of experience and interest in the topics covered in the Mathematics of the Mind course. n = 18 students.

We then asked the students questions related to their interest in the class and the topics discussed up to this point, and before the execution of live experiments. We found that 94.4% of the students reported that they are interested in the applications of mathematics to biological processes (Figure 3E), and all of them (100%) wanted to learn about Internet-of-Things technologies (Figure 3F). Similarly, 94.4% of the students reported that this course required them to think at a deep level or use critical thinking (Figure 3G). In addition, 83.3% of the students thought that this course allowed them to examine questions that matter beyond the classroom (Figure 3H). When asked about the biological side of the class, 77.7% of the students were interested in learning about stem cells (Figure 3I) and 94.4% of them were interested in learning about neuroscience (Figure 3J). Similarly, 88.8% of the students claimed to be interested in learning about “connectoids” (Figure 3K), long-range connected organoids (Adewole et al., 2021; Cullen et al., 2019; Kirihara et al., 2019). When asked about the technologies to be used in the experiments, 66.6% of the students were comfortable using WetAI (Figure 3L), our custom interface with the experiment (Baudin et al., 2022b), and 72.2% of the students claimed to be comfortable using Jupyter notebooks to perform experiments (Figure 3M). We found that 61.1% of the students said they were comfortable reading neuroscience literature (Figure 3N), while 44.4% of the students felt comfortable performing high density MEA experiments at this time (Figure 3O). In addition, 66.6% of the students reported that they would be comfortable identifying spike burst signals in high density MEA data (Figure 3P). Altogether, we observed that, consistent with their academic background, most students are more comfortable with the quantitative aspect of the class than with the biological section, although they demonstrate a high interest in learning topics in stem cell and neuroscience.

Given the qualities of these students, we then asked them questions related to their mathematics self-concept, as this metric has been previously shown to be a predictor of performance (Colmar et al., 2019; Cribbs et al., 2021; Rueda-Gómez, et al., 2023). To do so, we asked the students their level of agreement with different statements. We found that 61.1% of the students agreed with the statement “I am capable and skillful at Mathematics” (Figure 4A), while 44.4% agreed with the statement “Being a good mathematics student makes me feel that my classmates and teachers think more of me’” (Figure 4B). These results are similar to previous reports of mathematics self-concept in university students (Rueda-Gómez, et al., 2023). However, unlike previous reports, we find that half of the students (50%) agree with the statement “In the mathematics exams I feel unsure, desperate, and nervous” (Figure 4C), while the minority of the students (44.4%) agree with the statement “My performance in mathematics largely depends on the methodology and empathy with teachers” (Figure 4D). This self-concept profile is consistent with university students who are high achievers in online mathematics training (Rueda-Gómez, et al., 2023), suggesting that our students were likely to succeed in a remote PBL experiment. Finally, when asked whether they agreed with the statement “Mathematics is useful and necessary in all areas of life”, we found that 83.3% of the students agreed (Figure 4E), while 88.8% of the students agreed with the statement “Mathematics is useful and necessary for a career in Biology” (Figure 4F). The combination of the students’ quantitative skills and interests, as well as their self-concept consistent with high achievers in mathematics, allowed us to conclude that this cohort of students is ideal for implementing remote PBL experiments using high density MEAs.

**Figure 4.**
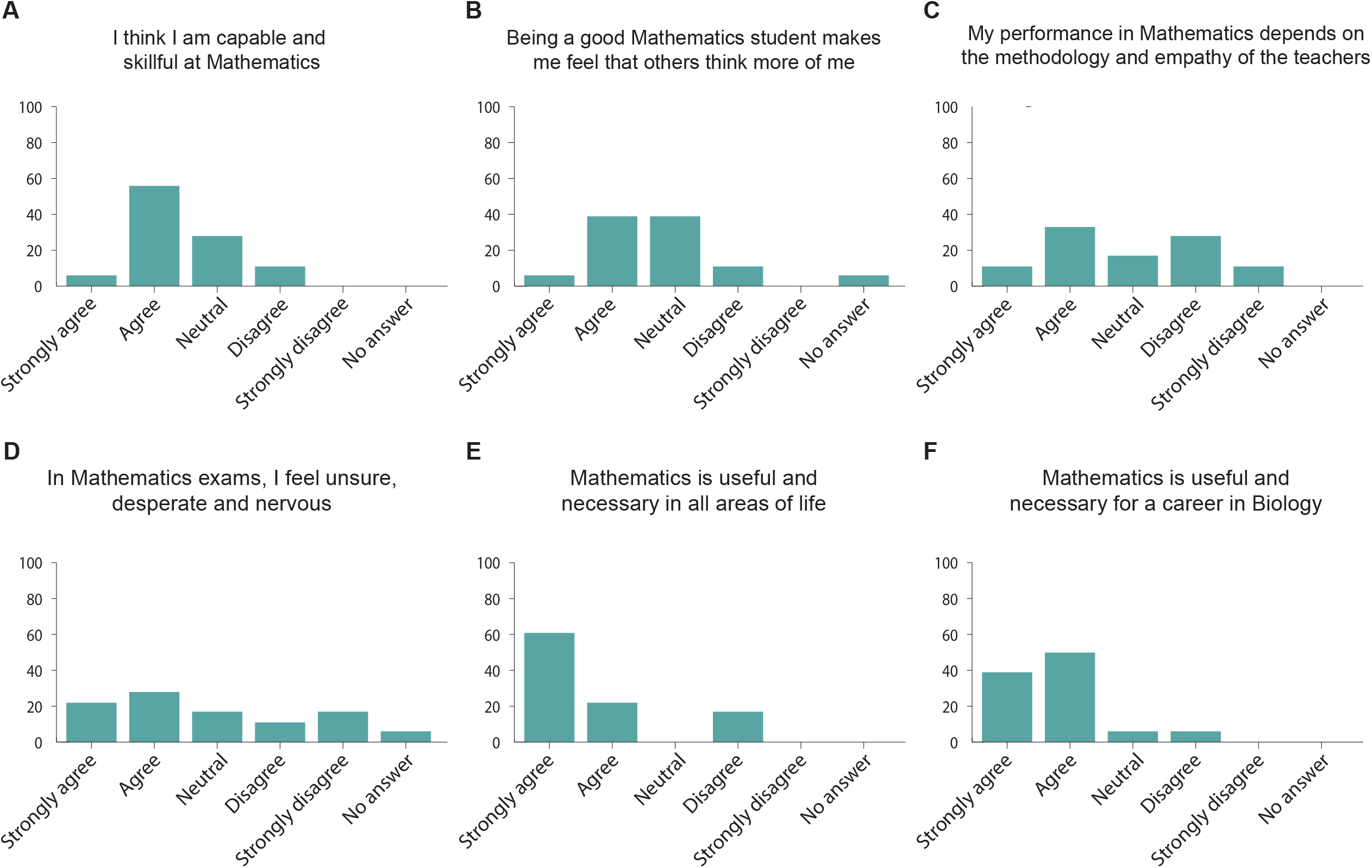
Mathematics self-concept of the students. A-F. Students’ answers to a survey assessing their mathematics self-concept. n = 18 students.

As part of the class, we asked students to work in groups to design and program stimulation patterns to be given to cortical organoids. They were asked to propose a hypothesis on what changes they expected to see in network behavior following stimulation. The students then analyzed the results of their stimulation patterns and presented their findings to the class. Each group was assigned two organoids: one to be used as an experimental organoid, and one control (Figures 5A-B). To help the students conduct their electrophysiology experiment, we designed a simple application programming interface (API). Each team of students had the opportunity to watch their stimulation experiment happen on the organoid live, in real time. The experiments were hosted via Zoom, with the class TA presenting the changes in neuronal activity that occurred while the organoids were being stimulated.

**Figure 5.**
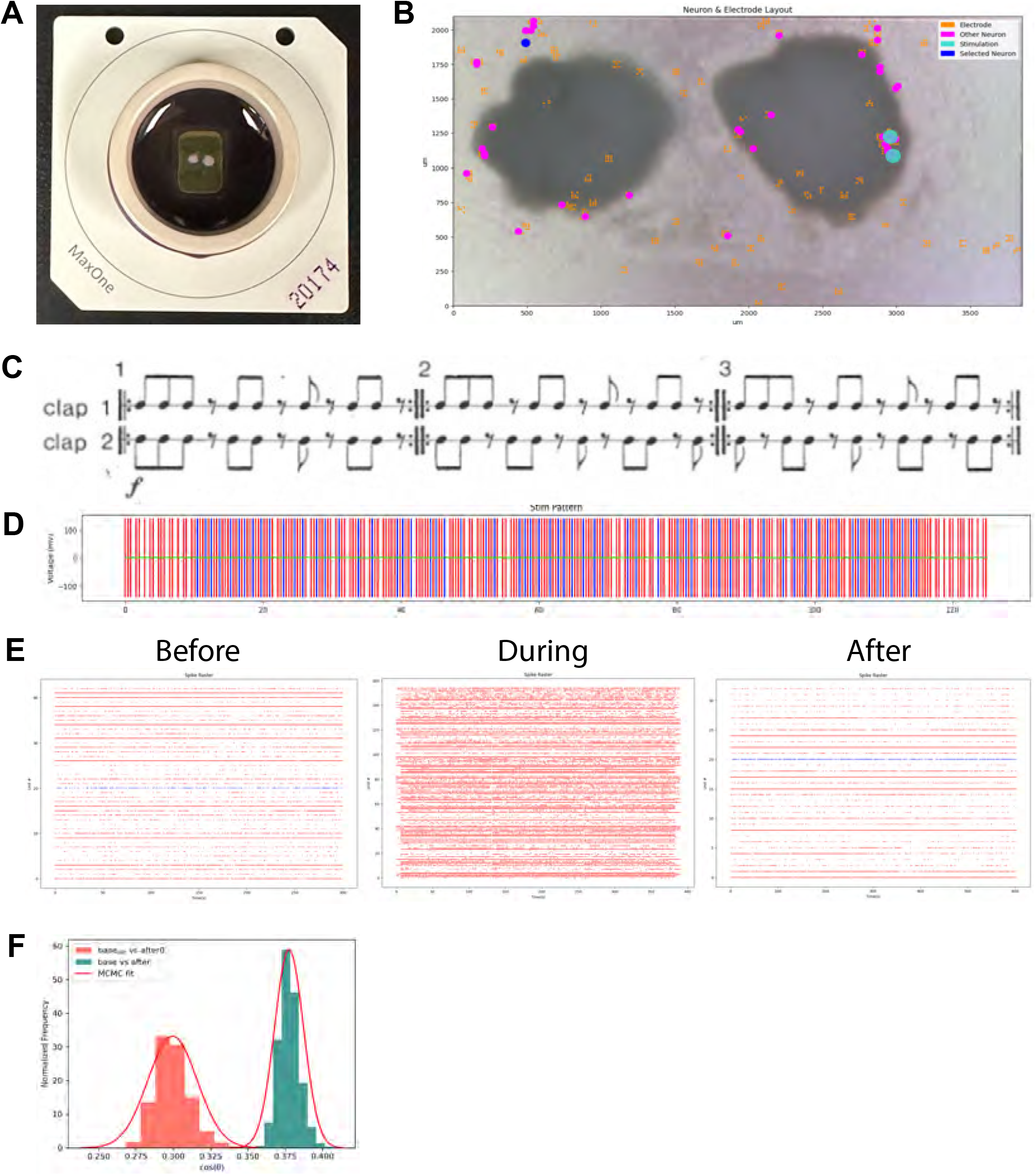
Student experiments explore programming neuronal circuits. A. Students were assigned two cerebral organoids (a test and control) to use in an experiment they designed. The organoids were placed on an HD MEA, which was used to send their electrophysiology stimulus pattern. B. The students used online software to explore the data. The software shows the location of neurons across the two organoids it is interacting with, as well as stimulation sites. Students can select a neuron to view specific information from it, such as the spike raster. C. The experiment shown involved a team of students who decided to code the rhythm of Steve Reich’s duet, Clapping Music. Each performers’ pattern was sent to different neurons. Image is from Reich’s music score. D. The students use the online software to verify that the stimulation pattern sent to the neurons follows the rhythm of the music. Red and blue represent the stimulation patterns sent to 2 different neuronal sites. E. The students were given the neuronal spike raster data from before, during and after the experiment. These graphs display the time points at which different neurons fire action potentials. Students can determine if the stimulus pattern they created evoked the predicted neuronal response by comparing the firing patterns between neurons. F. Example of the types of computational analysis students performed. To see if the stimulation had an effect on the neurons, this team performed a statistical test on the cosine similarity score, a metric of correlation. The test compared spontaneous activity directly after stimulation to that of 10 minutes after. They noticed that there was a statistically significant difference in the distribution of the cosine similarities between the two timepoints. MCMC fit = Markov-Chain Monte Carlo fit.

The API mentioned above provided students with considerable freedom in the types of experiment they could construct. One notable group, for example, translated the rhythm of a minimalist song: Steve Reich’s Clapping Music (Figures 5C-D). In this song, two artists clap at rhythms that synchronize and desynchronize throughout the song (Haack, 1991, 1998) (Figures 5C-D). The students wanted to see if this syncing of stimulation patterns might yield a Hebbian-like learning outcome. This rhythm has been previously used in mathematics problems in combinatorics and group theory (Haack, 1991, 1998). The students translated each of the clapping sequences to an electrode stimulation pattern using the API (Figures 5D). These patterns were then applied to the experimental organoid (Figure 5E) and the students found that the stimulation pattern had a measurable effect on neuronal behavior (Figures 5F).

The other groups created experiments of similar rigor and creativity (Supplemental Figure 1). Three of the five groups stimulated organoids with a high frequency tetanic pattern (Tamura et al., 2020). This experiment was popular because it was explained to have a high chance of prominent changes. Each group approached the problem using a unique protocol they designed. One group used the freedom that the stimulation API provided to design an experiment which generated a random walk over stimulation amplitudes, producing a unique sequence during each generation. This group was interested in whether these stimulation sequences produced unique patterns between neurons connected in a circuit.

The students were told to rigorously analyze the resulting data from their experiment to determine whether the experiment’s hypothesis was correct through techniques they were taught throughout the class: correlation matrix, spike time tiling coefficient, interspike intervals, and inter-neuronal spike latencies. Most importantly, the students had to implement some form of original analysis not taught during the class. This could either be a method learned from reading literature, or a novel technique they design themselves. Most groups implemented either a basic statistical test to verify their hypothesis, or an unsupervised machine learning technique to help interpret results. However, some groups developed novel analysis methods. Of note, one group implemented the underlying algorithm used in Principal Components Analysis (PCA) but using the spike time tiling matrix as the input into the algorithm (Supplemental Figure 1A-D). The group showed that this new technique was able to discern the test from control organoid based on neural signals (Supplemental Figure 1D).

### Organoid experiments lead to higher interest in stem cell and neuroscience research in mathematics students

To understand how the integration of HD MEA recording and stimulation of neuronal cell cultures in the classroom affects the students’ interest in neuroscience and stem cell research, we surveyed the Mathematics of the Mind students immediately after their presentations. All 24 students enrolled in the class responded to the survey. We found that 95.8% of the students said that they enjoyed learning about organoids (Figure 6A), and 91.6% enjoyed performing these experiments (Figure 6B). In addition, we found that 91.6% of the students thought that performing remote electrophysiology experiments was interesting (Figure 6C), while 95.8% of the students thought that the experiment selected helped solidify the concepts discussed in the class (Figure 6D).

**Figure 6.**
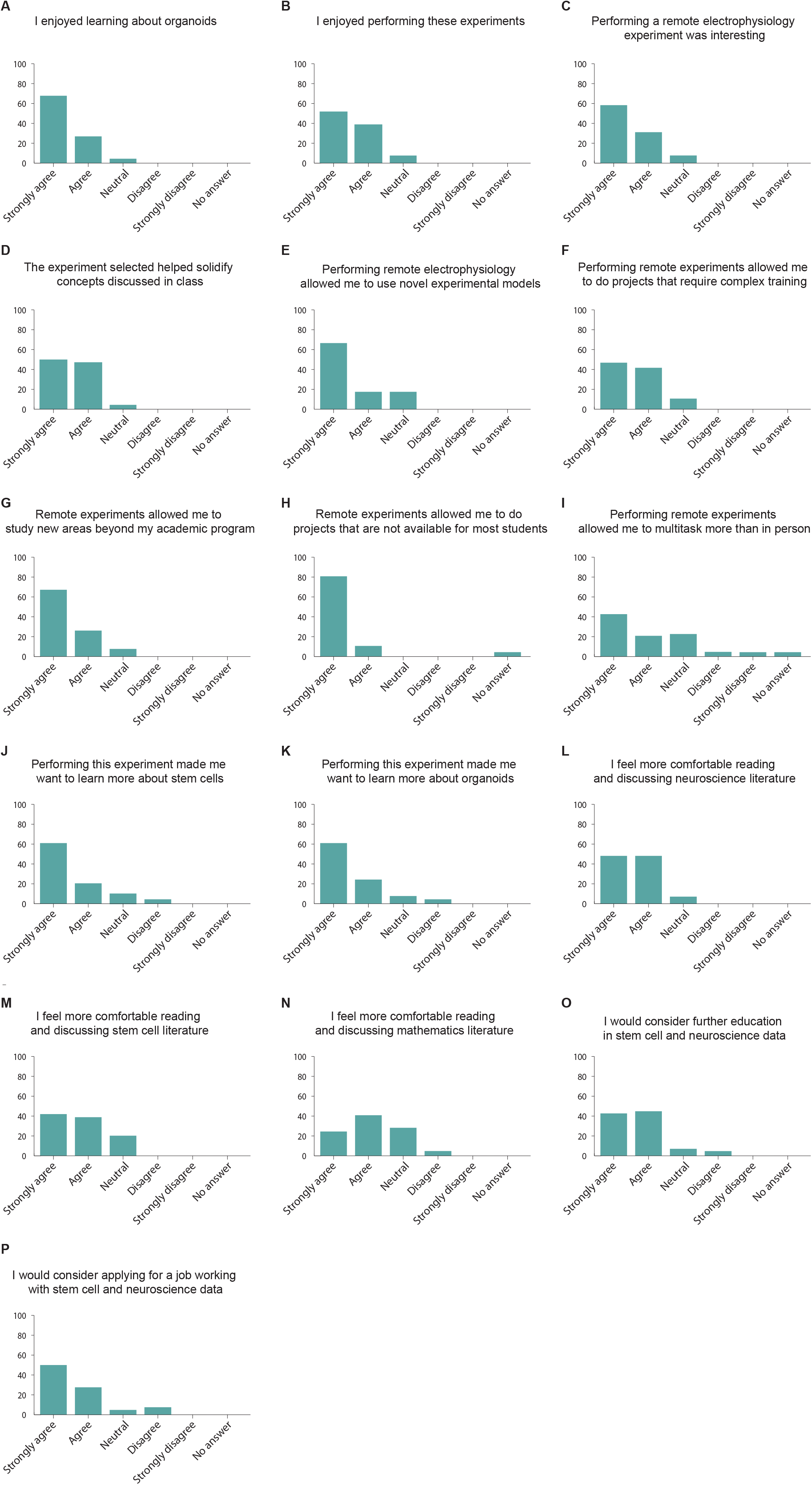
Mathematics students report positive attitudes toward stem cells and neuroscience topics after performing experiments with cortical organoids. (A-P) Post-experiment survey results for the Mathematics of the Mind course. n = 24 students.

We then asked the students for their perceptions after performing remote electrophysiology experiments. We found that 83.3% of the students thought that performing these experiments allowed them to perform novel and complex experiments (Figure 6E), while 87.5% of the students thought that this approach allowed them to do experiments that required complex training (Figure 6F). 91.6% of the students thought that these experiments allowed them to study new areas beyond their academic program (Figure 6G). 95.8% of the students considered that performing remote experiments allowed them to do projects that are not available to most students around the world (Figure 6H), while 62.5% of the students thought that performing remote experiments allowed them to multitask more than in person labs (Figure 6I).

Then, we focused on understanding how performing these experiments influenced the students’ interest in neuroscience and stem cell research. We found that 83.3% of the students thought that performing these experiments increased their interest in stem cells (Figure 6J) while 87.5% reported an increased interest in organoids (Figure 6K). Remarkably, we also find that after this course most of the students report a higher comfort in reading and discussing scientific literature in neuroscience (91.6%, Figure 6L), stem cells (79.1%, Figure 6M) and mathematics (66.6%, Figure 6N).

Finally, we asked the students how this work affected their career prospects. We found that 87.5% of the students would consider continuing further education in stem cells and neuroscience (Figure 6O) and 79.1% would consider applying to jobs analyzing stem cell and neuroscience data (Figure 6P). Altogether we conclude that our approach positively affected the students’ interest in stem cell and neuroscience.

## DISCUSSION

The impact of regenerative biology in the biotechnology and translational medicine sectors has grown exponentially over the past decades (Jacques et al., 2020). Therefore, there is an increased need for exposing students and young professionals to topics related to stem cells and regenerative medicine, including training in experimental design and applications of the technology (Sterner et al., 2020; W Sandoval et al., 2022; Wyles et al., 2019). Scalable approaches that can keep students motivated and engaged during their training should be prioritized to fulfill this demand (Wyles et al., 2020). The use of internet-enabled platforms can inexpensively scale the number of targeted students and reach traditionally underserved communities (Baudin et al., 2022b; Findyartini et al., 2021). Indeed, the use of virtual programs in the topic of regenerative medicine have been explored, especially during the COVID-19 pandemic (Wyles et al., 2020). Yet, in passive online teaching approaches maintaining students’ interest is challenging (Gares et al., 2020), and as a result, virtual courses often have a high dropout rate (Wang et al., 2023).

PBL approaches in STEM are effective at retaining students, especially those of underrepresented backgrounds (Ferreira et al., 2019). PBL courses are particularly successful when they integrate themes that are trendy in society (Carosso et al., 2019; Cotner and Gallup Jr, 2011). To this end, the high interest of the public and popular media in brain organoids (Presley et al., 2022) can make them ideal tools to train the next generation of students in regenerative medicine. Here, we took advantage of cortical organoids to design PBL modules and target students both in the life sciences, as well as other STEM disciplines, including mathematics, physics, engineering, and computer science. Both groups of students reported that performing the experiments in organoids helped solidify the concepts discussed in class. Furthermore, we showed that this approach increased the interest in topics related to neuroscience and stem cell research of both groups of students. In addition, non-biology students showed a higher interest in continuing their education in these topics. Moreover, the internet connectivity of our laboratory hardware allowed the students to perform the experiments remotely, up to 125 km from the experimental site. Additionally, this internet connectivity enables us to scale the number of users at a marginal price (Baudin et al., 2022b; Ly et al., 2021). These two properties open the possibility of massively deploying cloud-enabled technologies for training in neuroscience and regenerative biology.

We piloted this approach by adding a cortical organoid-based remote experiment module to two courses: a cell culture techniques and a mathematics course. However, organoids can be used in a variety of other PBL courses. For example, cortical organoids of multiple species (Mostajo-Radji et al., 2020; Pollen et al., 2019) can easily be integrated in an evolutionary biology course. “Assembloids”, fused organoids of different brain regions (Paşca, 2019), can be used to train students in systems neuroscience. Similarly, connectoids (Adewole et al., 2021; Cullen et al., 2019; Kirihara et al., 2019), can be used in the context of teaching higher level interactions between brain areas. Moreover, organoids can become important tools to teach chemistry in the context of drug screens (Salick et al., 2021; Schuster et al., 2020). Bioengineering courses can benefit from organoids experiments to teach biomaterials and sacrificial networks (Grebenyuk and Ranga, 2019). Finally, deriving organoids from multiple body regions beyond the brain, could be of benefit to physiology and pharmacology courses (Clevers, 2016).

In conclusion, we provided a proof of principle study using organoids as novel pedagogical tools for undergraduate education. We developed PBL-based curricula using imaging and electrophysiological tools. By connecting these technologies to the cloud, we were able to teach the courses remotely, while at the same time scaling to allow multiple students to interact with the same experimental platform. This approach led to a higher interest of the students toward stem cell and neuroscience paths. Altogether, this approach has the potential to greatly expand training in regenerative medicine and neuroscience and reach students currently underserved in their communities.

## EXPERIMENTAL PROCEDURES

### Ethics statement

The UCSC Institutional Review Board (IRB) reviewed this work at the proposal stage and determined that it did not constitute human subject study. Therefore, the work was IRB exempt.

### Students

The students were part of two different courses: 1) The students who performed the remote microscopy experiments were undergraduate students enrolled in the Techniques in Tissue Culture course at the University of San Francisco. A total of 10 students were enrolled in this course.

2) The students who performed the electrophysiology experiments were enrolled in the BME118: Mathematics of the Mind course at the University of California Santa Cruz. A total of 24 students were enrolled in this course.

### Surveys

All surveys were performed using Google Forms. The students received the link to the survey from their instructors. All surveys were completely anonymous and we did not record any identifiable information from any of the students. All students were informed that the answers to the surveys were anonymous and would not influence their grades.

For the Techniques in Tissue Culture course, we performed one survey at the end of the course. All 10 students who were part of the course responded to this survey. For the Mathematics of the Mind course, we performed 2 surveys: one before the experiment, and one after the experiment. For the pre-experiment survey, 18 students responded. For the post-experiment survey, all 24 students responded.

### Embryonic stem cell culture

All experiments were performed in the ES-E14TG2a mouse embryonic stem cell (ESC) line (ATCC CRL-1821). This line is derived from a male of the 129/Ola mouse strain. Mycoplasma testing confirmed lack of contamination.

ESCs were maintained on Recombinant Human Protein Vitronectin (Thermo Fisher Scientific # A14700) coated plates using mESC maintenance media containing Glasgow Minimum Essential Medium (Thermo Fisher Scientific # 11710035), Embryonic Stem Cell-Qualified Fetal Bovine Serum (Thermo Fisher Scientific # 10439001), 0.1 mM MEM Non-Essential Amino Acids (Thermo Fisher Scientific # 11140050), 1 mM Sodium Pyruvate (Millipore Sigma # S8636), 2 mM Glutamax supplement (Thermo Fisher Scientific # 35050061), 0.1 mM 2-Mercaptoethanol (Millipore Sigma # M3148), and 0.05 mg/ml Primocin (Invitrogen # ant-pm-05). mESC maintenance media was supplemented with 1,000 units/mL of Recombinant Mouse Leukemia Inhibitory Factor (Millipore Sigma # ESG1107), 1 µm PD0325901 (Stem Cell Technologies # 72182), and 3 µm CHIR99021 (Stem Cell Technologies # 72054). Media was changed every 2-3 days.

Vitronectin coating was incubated for 15 min at a concentration of 0.5 µg/mL dissolved in 1X Phosphate-buffered saline (PBS) pH 7.4 (Thermo Fisher Scientific # 70011044). Dissociation and cell passages were done using ReLeSR passaging reagent (Stem Cell Technologies # 05872) according to the manufacturer’s instructions. Cell freezing was done in mFreSR cryopreservation medium (Stem Cell Technologies # 05855) according to the manufacturer’s instructions.

### Cortical organoids generation

Mouse cortical organoids were grown as previously described by our group (Park et al., 2023). To generate cortical organoids we clump-dissociated ESCs using ReLeSR and re-aggregated in lipidure-coated 96-well V-bottom plates at a density of 5,000 cells per aggregate, in 150 µL of mESC maintenance media supplemented with Rho Kinase Inhibitor (Y-27632, 10 µM, Tocris # 1254), 1 µm PD0325901 (Stem Cell Technologies # 72182), 3 µm CHIR99021 (Stem Cell Technologies # 72054) (Day -1).

After one day (Day 0), we replaced the medium with cortical differentiation medium containing Glasgow Minimum Essential Medium (Thermo Fisher Scientific # 11710035), 10% Knockout Serum Replacement (Thermo Fisher Scientific # 10828028), 0.1 mM MEM Non-Essential Amino Acids (Thermo Fisher Scientific # 11140050), 1 mM Sodium Pyruvate (Millipore Sigma # S8636), 2 mM Glutamax supplement (Thermo Fisher Scientific # 35050061) 0.1 mM 2- Mercaptoethanol (Millipore Sigma # M3148) and 0.05 mg/ml Primocin (Invitrogen # ant-pm-05). Cortical differentiation medium was supplemented with Rho Kinase Inhibitor (Y-27632, 20 µM # 1254), WNT inhibitor (IWR1-ε, 3 µM, Cayman Chemical # 13659) and TGF-Beta inhibitor (SB431542, Tocris # 1614, 5 μM, days 0-7). Media was changed every other day until day 7.

On day 7 organoids were transferred to ultra-low adhesion plates (Millipore Sigma # CLS3471) where media was aspirated and replaced with fresh neuronal differentiation media. The plate with organoids was put on an orbital shaker at 60 revolutions per minute. Neuronal differentiation medium contained Dulbecco’s Modified Eagle Medium: Nutrient Mixture F-12 with GlutaMAX supplement (Thermo Fisher Scientific # 10565018), 1X N-2 Supplement (Thermo Fisher Scientific # 17502048), 1X Chemically Defined Lipid Concentrate (Thermo Fisher Scientific # 11905031) and 0.05 mg/ml Primocin (Invitrogen # ant-pm-05). Organoids were grown under 5% CO_2_ conditions. Medium was changed every 2-3 days.

On day 14 and onward, we added 5 µg/mL Heparin sodium salt from porcine intestinal mucosa (Millipore Sigma # H3149) and 0.5% v/v Matrigel Growth Factor Reduced (GFR) Basement Membrane Matrix, LDEV-free (Matrigel GFR, Corning # 354230) to the neuronal differentiation medium.

On day 21 and onward, we transferred the organoids to neuronal maturation media containing BrainPhys Neuronal Medium (Stem Cell Technologies # 05790), 1X N-2 Supplement, 1X Chemically Defined Lipid Concentrate (Thermo Fisher Scientific # 11905031), 1X B-27 Supplement (Thermo Fisher Scientific # 17504044), 0.05 mg/ml Primocin (Invitrogen # ant-pm-05) and 0.5% v/v Matrigel Growth Factor Reduced (GFR) Basement Membrane Matrix, LDEV-free.

### Microscopy experiments

The mouse cortical organoids were the basis of microscopy experiments. The microscope used to image these organoids is a variant of a previously described microscope (Baudin et al., 2022), named the Streamscope. The Streamscope is a low cost and open hardware 6-well microscope designed to perform simultaneous longitudinal imaging of cell culture in six separate wells. It produces timelapse sequences tracking morphological changes in cell cultures. The device is constructed from a combination of off-the-shelf and custom-made components. The off-the-shelf components include a motor driver (Pololu Tic T825), a motor linear actuator module, an LED light panel, and 6 USB cameras (HBVCAM 5MP module with Omnivision 5640 sensor.) The rest of the microscope is composed of parts which can be 3D printed or CNC milled in-house on consumer equipment or ordered from a third-party manufacturing service.

The organoids were transferred and adhered to a 6 well plate (6 well Nunc™ Cell-Culture Treated Multidishes, Thermo Fisher Scientific #140685). Adhesion was accomplished by transferring 0.5 mL of Matrigel Growth Factor Reduced (GFR) Basement Membrane Matrix, LDEV-free (Matrigel GFR, Corning # 354230) into a 1.5 mL Eppendorf Safe-Lock Tube (Thermo Fisher Scientific #22363204) kept on ice. The organoid was transferred into the matrigel with a wide bore pipette, pipetting up and down gently 5 times to ensure maximal coating. The matrigel coated organoid was then transferred to the 6 well plate and allowed to adhere for 10 minutes at 37° C before adding in fresh neuronal maturation media. For the experiments, we tested the effects of two drugs: 1µM of the proapoptotic drug staurosporine (Thermo Fisher Scientific # NC1401148) and 1µM of the DNA topoisomerase type I inhibitor camptothecin (Thermo Fisher Scientific # 501361118). In addition, the students had control organoids in which no drugs were added.

Two plates of organoids atop two Streamscopes were used for this experiment, totalling twelve potential wells worth of images. Images of each well were taken approximately every 60 seconds over a period of 5 days while the cultures were in the incubator. The images were streamed through YouTube, allowing the users to access the data in real time, as well as after the end of the experiment.

After the experiment was terminated, images from each condition were turned into a timelapse. The images for each well were converted into timelapses using Adobe Premiere. Images were computer analyzed for features such as organoid area.

### Organoid area measurements

To reduce the computational load on ImageJ, the software used to calculate the area of the organoids, the time lapse images were subsampled into sparser timelapse images. Starting with the 0-hour time point, images of organoids were sampled every 6 hours for 72 hours total. By interpolating, adjusting threshold parameters in ImageJ and using the tracing tool, areas were determined for each organoid at all time points.

### Electrophysiology experiments

We plated mouse cortical organoids at day 22 on MaxOne high density multielectrode arrays (Maxwell Biosystems # PSM). Prior to organoid plating, the multielectrode arrays were coated in 2 steps: First, we performed an overnight coating with 0.01% Poly-L-ornithine (Millipore Sigma # P4957) at 37° C overnight. Then washed the plates 3 times with PBS. We then performed an overnight coating with 5 µg/ml mouse Laminin (Fisher Scientific # CB40232) and 5 µg/ml human Fibronectin (Fisher Scientific # CB40008) at 37° C.

After coating, we placed the organoids on the chip and removed excess media. The organoids were then incubated at 37° C for 20 minutes to promote attachment. We then added prewarmed BrainPhys Neuronal Medium supplemented with 20 ng/mL recombinant human brain-derived neurotrophic factor (Stem Cell Technologies # 78005), 1X N-2 Supplement (Thermo Fisher Scientific # 17502048), 1X Chemically Defined Lipid Concentrate (Thermo Fisher Scientific # 11905031), 1X B-27 Supplement (Thermo Fisher Scientific # 17504044), 0.05 mg/ml Primocin (Invitrogen # ant-pm-05) and 0.5% v/v Matrigel Growth Factor Reduced (GFR) Basement Membrane Matrix, LDEV-free. We changed the media every 2-3 days.

Using the Maxwell Biosystems software, activity scans were performed every 2-3 days. We sampled signals from 1024 of the ∼26,000 electrodes at a time in a sweeping formation across the MEA at an interval of 30-45 seconds for each configuration. A base level of activity was measured by electrical events which crossed the 5-root mean square (rms) noise threshold. If activity was detected, 5–12-minute recordings were taken using Maxwell Biosystems’ electrode selection algorithm which maximizes clusters of electrodes around hotspots of high relative firing rate. Neurons were identified by analyzing the neuronal footprint – the signal averaged waveform across a patch of multiple electrodes surrounding the potential neuron. Neurons chosen to be stimulated required detection of at least 10 threshold-crossing events. We chose stimulation channels as the channels that recorded the highest amplitude signals from the neuron to have the highest probability of evoking action potentials (Radivojevic et al., 2016).

### Data processing

Raw electrical data from the MEA was saved during each of the experiments in the hdf5 format. This data was spike-sorted into individual units (putative neurons) using Kilosort 2 (Pachitariu et al., 2023). This process filters the data, whitens (Pachitariu et al., 2023), then clusters neurons based on matching spike-waveform templates. For each identified neuron, this process outputs a list of spikes times together with the spatial location on the MEA. To validate the biological plausibility of the identified neurons, expert human curation was carried out through the open-source software Phy (https://github.com/cortex-lab/phy). Putative neural units are retained if they have key biological features such as a biologically consistent shape and duration for their action potential waveforms. Each experiment recording was individually processed with the same set of spike sorting parameters. Electric stimulation artifacts and noise were removed during the curation. This data was delivered to the students in Python Numpy format allowing easy subsampling.

### Electrophysiology Experimental Design

Mathematics of the Mind students were introduced to the idea of analyzing electrophysiological measurements by conducting analysis from a previous set of published electrophysiological recordings in human organoids (Sharf et al., 2022). The students were taught to utilize measures including spike-time tiling (Cutts and Eglen, 2014), correlation, inter-spike-intervals, and latency distributions to characterize neural circuit behavior. After this initial homework assignment, students were asked to propose a circuit perturbation experiment using electrode-supplied stimulation on live tissue to augment the underlying neural circuitry, where they would analyze spontaneous activity before and after the perturbation. The setup of the experiment involved a five-minute baseline recording, a five minute recording with student-designed stimulation, and a fifteen minute post-stimulation recording. The students were specifically tasked with designing and coding their own stimulation paradigm to be executed during their experiment, and proposing a hypothesis for changes that would occur between the baseline and post-stimulation recordings. They were encouraged to be creative.

### Stimulation Programming

To make the task of stimulation paradigm design feasible and straightforward for the Mathematics of the Mind students, a simple application programming interface (API) was designed. The students were told that they could stimulate three neurons in any manner that the API enabled, and they could select the approximate distance of the three neurons as a proxy for connectivity. At a high level, the students craft sequences of stimulations using the API, and choose a frequency to iterate through these stimulation sequences. The API enables the creation of stimulation paradigms by three simple commands: ‘stim’, ‘delay’ and ‘next’. The ‘stim’ command has three parameters: 1) the list of neurons to stimulate, b) the amplitude in millivolts to stimulate, 3) the width of the biphasic pulse. Recommended values of 150 mV for the amplitude and 200 microseconds per phase for the biphasic pulse were given to students. The ‘delay’ command has one parameter: the time to delay in milliseconds. It pauses the stimulation. The ‘next’ command effectively ends the current stimulation sequence.

## ACKNOWLEDGMENTS

We would like to thank the students of University of San Francisco and the University of California Santa Cruz for their enthusiasm and dedication during this project. We would also like to thank Ryan Hoffman and Claudia Paz Flores for providing support during the Mathematics of the mind course. In addition, we are thankful to Sri Kurniawan, Catharina Lindley and Sofie Salama for their critical feedback on this manuscript and support during the execution of this project.

This work was supported by Schmidt Futures (SF857) to M.T. and D.H.; National Human Genome Research Institute (1RM1HG011543) to M.T. and D.H.; National Science Foundation (NSF2134955) to M.T. and D.H. (NSF2034037) to M.T.; the National Institute of Mental Health (1U24MH132628) to D.H. and M.A.M.-R. H.E.S. is a National Science Foundation Graduate Student Research Fellowship grantee. K.V. was supported by the ARCS Foundation and grant T32HG012344 from the National Human Genome Research Institute (NHGRI). J.L.S. was supported by the University of California Santa Cruz Chancellor’s Postdoctoral Fellowship, the NIH K12GM139185 (through NIGMS to UCSC IBSC), and LRP0000018281 (NICHD). We are thankful to the Pacific Research Platform, supported by the National Science Foundation under Award Numbers CNS-1730158, ACI-1540112, ACI-1541349, OAC-1826967, the University of California Office of the President, and the University of California San Diego’s California Institute for Telecommunications and Information Technology/Qualcomm Institute. Thanks to CENIC for the 100 Gbps networks.

## DATA AVAILABILITY STATEMENT

The consent given by the participants does not allow for open storage of data on an individual level in public repositories. Anonymized survey data are available upon request by qualified scientists. Requests require a concept paper describing the purpose of data access, ethical approval at the applicant’s institution, and provision for secure data access.

## AUTHOR CONTRIBUTION STATEMENT

Performed the experiments: M.A.T.E., H.E.S., A.R., S.V-C., D.E., K.V., J.G.

Analyzed and interpreted the data: A.R., S.H., S.V.-C., J.G.

Contributed reagents, materials, analysis tools or data: M.A.T.E., H.E.S., A.R., S.V.-C., D.E., K.V., J.G., J.L.S., Y.R.M., M.T.

Wrote the paper: M.A.T.E., H.E.S., A.R., M.A.M.-R.

Conceived and designed the experiments: N.O.W., D.H., M.A.M.-R.

