## Supplemental Information for "Internet-connected cortical organoids for project-based stem cell and neuroscience education"

Biorxiv
Supplemental Information

### Internet-connected cortical organoids for project-based stem cell and neuroscience education

Matthew A.T. Elliott*, Hunter E. Schweiger*, Ash Robbins*, Samira Vera-Choqqueccota, Drew Ehrlich, Sebastian Hernandez, Kateryna Voitiuk, Jinghui Geng, Jess L. Sevetson, Yohei M. Rosen, Mircea Teodorescu, Nico O. Wagner, David Haussler, Mohammed A. Mostajo-Radji

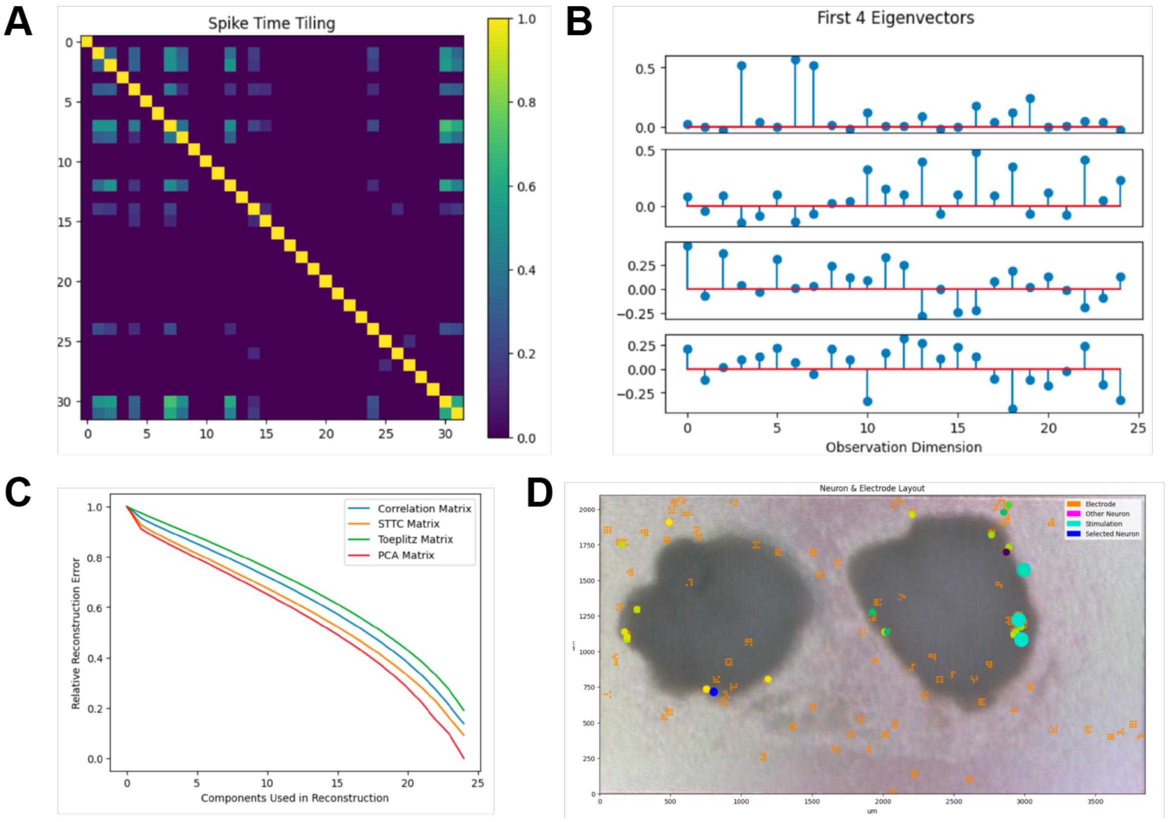


**Supplemental Figure 1. Example of Mathematics of the mind student-project applying tetanic pattern to cortical organoids.** A. Spike time tilling matrix for the electrophysiology data recorded from cortical organoids. B. Eigendecomposition on the spike time tilling matrix to extract its eigenvalues and eigenvectors. For illustration, the first 4 eigenvectors are plotted. C. Comparison of eigendecomposition on the spike time tiling matrix to other popular correlation techniques. Specifically, they considered how the method's reconstruction error compared to that of performing principal components analysis on the correlation matrix. D. Superimposition of the first eigenvector's values (represented as small circles in the blue-yellow range) on top of their corresponding neural unit allowed the students to differentiate which neurons came from which organoid.
